# Unraveling the Self-Assembly Mechanisms in Bacterial Cellulose Hydrogels

**DOI:** 10.64898/2026.01.21.700946

**Authors:** Go Takayama, Tetsuo Kondo

**Author notes:** Corresponding author: Tetsuo Kondo. (G. Takayama).

## Abstract

Bacterial cellulose (BC) hydrogels produced by *Gluconacetobacter* species hold considerable promise for a wide range of applications owing to their exceptional mechanical properties, biocompatibility, and biodegradability. Achieving precise control over their structural and mechanical characteristics is crucial for the engineering of BC-based materials. In this study, we investigated the formation dynamics and structural features of BC hydrogels, emphasizing the complex interplay between cellulose nanofibril secretion and bacterial motility. Comprehensive tracking of bacterial movement during hydrogel formation has validated mechanisms underlying the development of branching and merging junctions, which are key elements that define the network’s physical properties. Additionally, we observed the emergence of vortex-lattice and chiral-nematic structures during hydrogel development, depending on bacterial and cellulose densities. These insights contribute to a fundamental understanding of bottom-up 3D fabrication of BC hydrogels that harness the collective behavior of cellulose-producing bacteria.

## INTRODUCTION

Microorganisms belonging to the genus *Gluconacetobacter* are well known to produce cellulose in the form of gel-like membranes^1^. These bacteria secrete extracellular nanofibrils, referred to as “BC ribbons”^2^, which self-assemble into a network through physical cross-linking^3^, ultimately forming gel-like structures at the air–liquid interface of their culture medium. These BC hydrogels possess remarkable mechanical properties^4–7^, biocompatibility^8^, and biodegradability^9,10^, which derive from the structural features of BC nanofibrils (molecular and crystalline structures) and the hierarchical structure they form (network architecture^3,11,12^ and macroscopic morphology^12^). Such inherent properties have motivated extensive research into BC hydrogels for diverse applications, including medical devices^13^, food products^14,15^, cosmetics^16^, and textiles^17–19^. Recently, interest has revived in utilizing biological approaches to design BC-based materials^20–22^, particularly by leveraging *Gluconacetobacter*’s ability to secrete cellulose nanofibrils and construct complex macroscopic structures, thereby positioning these bacteria as promising scaffolds for responsive and adaptive “living materials”^20,23,24^.

One of the primary challenges in utilizing BC hydrogels is the variability in their physical properties. Cultivation conditions and bacterial strain selection can significantly influence the hydrogel characteristics, resulting in inconsistent properties^25–28^. Compared to dried BC films, which exhibit high elastic moduli (5–8 GPa^28^), BC hydrogels show much lower absolute stiffness. However, their mechanical properties span a broad range (elastic modulus: 4.3 ± 3.5 MPa^7,26,29^), suggesting the potential for a wide tunable range. Previous studies suggest that these variations originate from their network structures formed by *Gluconacetobacter*^3,27^. To realize diverse applications, it is essential to understand and control the structural formation processes that govern the mechanical and physical properties of BC hydrogels.

Understanding these processes requires insight into how bacterial cellulose secretion and cell motility contribute to the network formation. *Gluconacetobacter* bacteria propel themselves by exuding cellulose nanofibrils^2,30,31^—a unique motility mechanism distinct from flagellar-based movement seen in *Escherichia coli*^32^. The final BC hydrogel architecture reflects the dynamics of cellulose secretion coupled with bacterial movement during proliferation. Importantly, the assembly of higher-order structures in BC hydrogels—such as branched and interconnected networks—is not merely the result of fiber accumulation, but is tightly linked to the bacterial ecosystem dynamics, including nutrient consumption, proliferation, and nanofiber secretion activity. Thus, the BC hydrogel serves a physical record of bacterial growth, motility, and fiber secretion activities.

In this perspective, previous work has explored how interactions between cellulose fibers and substrates can guide the bacterial movement to produce patterns or aligned fiber structures, such as honeycomb formations^30,33–36^. Video microscopy studies have further elucidated the distinct secretion behaviors of *Gluconacetobacter,* including the frequent deposition of newly formed fibers onto existing BC ribbons, which reinforces the network and contributes to hydrogel robustness^30,33,34^. While most studies have focused on the motility of individual bacteria, the collective behavior of bacterial populations as they construct the fiber networks remains less understood. Since the macroscopic properties of BC hydrogels are intimately linked to network architecture, deciphering how bacterial communities influence fiber assembly, especially the formation of cross-linked structures, is critical. Controlling the density and arrangement of crosslink points could enable tuning of key material properties such as elasticity, strength, and extensibility.

In this study, we investigated the structural characteristics and formation dynamics of BC hydrogels produced by *Gluconacetobacter*. We characterized both isotropic fiber networks and more complex patterns such as chiral nematic and vortex structures. In situ observations of the movement of bacterial populations during hydrogel formation provided direct insights into mechanisms underlying the development of crosslinked structures that substantially impact the physical properties of BC hydrogels. We also observed the emergence of anisotropic structures as the bacterial and cellulose volume fractions increased. These findings establish a foundation for designing novel BC materials through manipulation of bacterial motility and activity, enabling advances in bottom-up 3D gel fabrication.

## EXPERIMENTAL SECTION

### Materials

The bacterial strains *Gluconacetobacter hansenii* ATCC53582 and *G. hansenii* ATCC23769 were obtained from the American Type Culture Collection (VA, USA) and preserved at −80°C. Bacto™ peptone and Bacto™ yeast extract were purchased from Difco Laboratories Inc. (Franklin Lakes, New Jersey, USA). Glucose, cellulase (Celluclast 1.5 L), and Fluorescent Brightener 28 (FB28; also known as Calcofluor White M2R) were acquired from Sigma-Aldrich (St. Louis, Missouri, USA). Disodium hydrogen phosphate heptahydrate (Na_2_HPO_4_·7H_2_O) was purchased from Nacalai Tesque Inc. (Kyoto, Japan). Citric acid was obtained from Wako Pure Chemical Inc. (Osaka, Japan). Silpot 184 was acquired from Dow Toray Co., Ltd (Tokyo, Japan).

### Bacterial culture

Bacterial cultures stored at −80°C were thawed and streaked onto Hestrin-Schramm (HS) agar^37^ plates containing 1% (w/v) agar. The culture plates were incubated at 30°C. For routine cultivation, bacterial suspensions squeezed from the BC hydrogels formed during static culture were transferred into fresh HS medium in 24-well plates and incubated at 30°C. These cultures were used as the inoculum for subsequent experiments.

### Measurements

#### Confocal Microscopic Observation of BC hydrogels

Confocal laser scanning microscopy (CLSM) was employed to analyze the microstructure of BC hydrogels. BC was produced by inoculating *Gluconacetobacter* into a liquid culture medium and incubating them at 30°C for several days. The resulting hydrogels were purified by immersion in a 0.1% (w/v) NaOH solution, heated at 80°C for 4 hours, then thoroughly rinsed with water. For visualization, the purified samples were stained with FB28 following established procedures^3^. After washing, samples were cut into approximately 1 cm sections and mounted on 35 mm glass-bottom dishes. Imaging was performed using a Leica TCS SP8 confocal microscope (Wetzlar, Germany) equipped with a 100× objective lens, under conditions described in previous studies^3^.

#### Time-lapse microscopic observation of BC hydrogel formation

To visualize the dynamics of BC hydrogel formation, time-lapse microscopy was conducted. Bacterial cultures were grown in 50 mL plastic tubes containing 5 mL of HS medium with 0.1% (v/v) cellulase (Celluclast 1.5L, Sigma Aldrich) to mitigate cellulose aggregation under continuous agitation (150 rpm) at 30°C. After incubation, the cultures were centrifuged at 3600 rpm for 3 minutes, and the pellets were then collected and resuspended in fresh HS medium without cellulase. A 10 μL aliquot of the suspension was introduced into a custom-made observation chamber (**Figure 1**). The chamber was made of oxygen-permeable polydimethylsiloxane (PDMS; Silpot 184) and contained a circular recess approximately 200 μm deep and 6 mm in diameter. It was securely mounted on a 35 mm glass-bottom dish containing HS agar medium solidified with 1% agar, allowing oxygen diffusion through PDMS and nutrient exchange from the agar medium solidified with 1% agar. Time-lapse images were captured at 12-second intervals using a phase-contrast microscope (CKX41, Olympus, Tokyo, Japan), equipped with either a CCD camera (Spot insight, SPOT Imaging, Michigan, USA) or a CMOS camera (MU1000-HS, AmScope, Irvine, California, USA) using a 40× objective lens (LCAch N PhP, Olympus, NA = 0.55).

**Figure 1.**
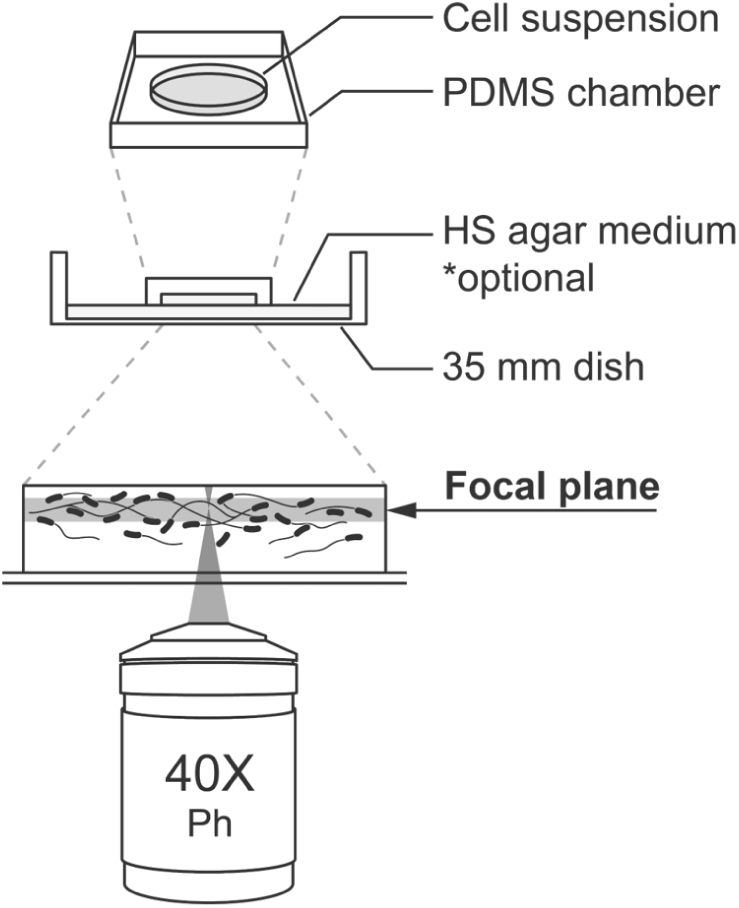
Schematic illustration of the experimental setup for time-lapse microscopy. The oxygen-permeable PDMS chamber enables stable live imaging of bacterial movement, with oxygen supplied through the PDMS and nutrients via HS agar.

#### Tracking analysis of bacterial movements

Bacterial motility during hydrogel formation was analyzed by tracking of time-lapse image sequences. Image pre-processing involved resizing, noise reduction using a rolling ball filter, and Gaussian blurring, performed with ImageJ FIJI^38^. Bacterial segmentation was achieved through intensity thresholding and shape fitting tailored for rod-shaped cells, utilizing plugins such as microbeJ^39^ and TrackMate^40^. Trajectories were reconstructed by linking bacterial positions frame by frame using a nearest-neighbor tracking. From these data, parameters such as instantaneous speed and the mean squared displacement (MSD) were calculated to quantify motility patterns.

#### Atomic Force Microscopy (AFM) observation of BC networks

To examine the nanostructure of the cellulose fiber network beyond the CLSM resolution, AFM imaging was performed. Glass substrates coated with poly-L-lysine were used to enhance bacterial cell and fiber adhesion. Dispersions of *Gluconacetobacter* in HS medium were drop-cast onto the substrates and incubated at 30°C for 30 minutes to promote cellulose fiber production. After rinsing with water and drying, samples were imaged with an MFP-3D-SA AFM system (Oxford Instruments, Santa Barbara, California, USA) equipped with a OMCL-AC240TS cantilever (spring constant = 2 N/m; Olympus Co., Tokyo, Japan).

#### Orientation Analysis

The orientation of bacterial cells in phase contrast images was quantified using the OrientationJ plugin for ImageJ^41^. This method applies structure tensor analysis to determine local orientation angle and coherency metrics, where coherence ranges from 0 (no preferred orientation) to 1 (highly aligned). This analysis provides quantitative insight into the spatial organization of the bacterial populations.

## RESULTS AND DISCUSSION

### Structural patterns in BC hydrogels visualized by confocal laser scanning microscopy

Confocal laser scanning microscopy (CLSM) analysis of BC hydrogels revealed occasional presence of two distinctive structural patterns (**Figure 2a–c**), which differ from the commonly observed isotropic fiber network architecture (**Figure 2a**). Specifically, **Figure 2b** illustrates a pattern characterized by interconnected domains exhibiting nematic alignment, while **Figure 2c** displays a vortex structure, in which fibers are arranged in a circular configuration. Although these unique morphologies are relatively rare, the latter two structures are anticipated to impart novel material properties distinct from those of the typical isotropic fiber network. Elucidating the mechanisms underlying the formation of these patterns is crucial for understanding the variability in the physical properties of BC hydrogels and for guiding the rational design of these materials.

**Figure 2.**
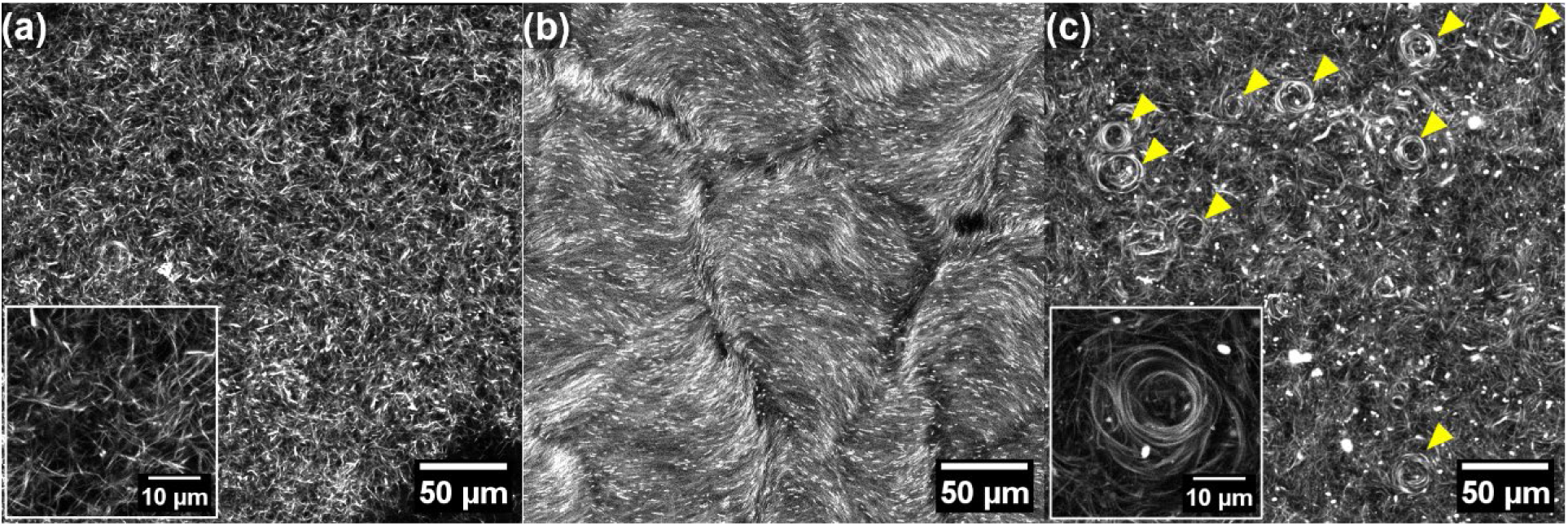
CLSM images of various BC pellicles showing different structural architectures: (a) isotropic fiber network (the most common), (b) anisotropic domains with nematic alignment, and (c) vortex structures. Arrowheads in (c) indicate vortex patterns.

### Time-lapse observation of BC hydrogel formation

The motility of *Gluconacetobacter* spp. plays a pivotal role in the development of the fibrous network within BC hydrogels. As obligate aerobes, these bacteria proliferate vigorously in oxygen-rich regions near the air–liquid interface during culture. Concurrently, they secrete cellulose ribbons, stacking into multiple two-dimensional layers that build up a three-dimensional hydrogel. To elucidate the dynamics of this process, time-lapse microscopy was performed using a custom-designed observation chamber (**Figure 1**), enabling stable imaging of bacterial movements within the liquid medium.

Analysis of the recorded time-lapse sequences (**Figure 3a**) involved color-coding bacterial positions according to elapsed time and superimposing them to generate a time-projection image (**Figure 3b**). By tracing the trajectories of individual bacteria across the sequences, the formation pathways of cellulose fibers—otherwise invisible in single-channel images—were implicitly visualized (**Figure 3b, c**). Notably, bacteria, which were not engaged in cellulose secretion, exhibited Brownian motion, diffusing randomly within the medium, thereby distinguishing them from actively secreting bacteria.

**Figure 3.**
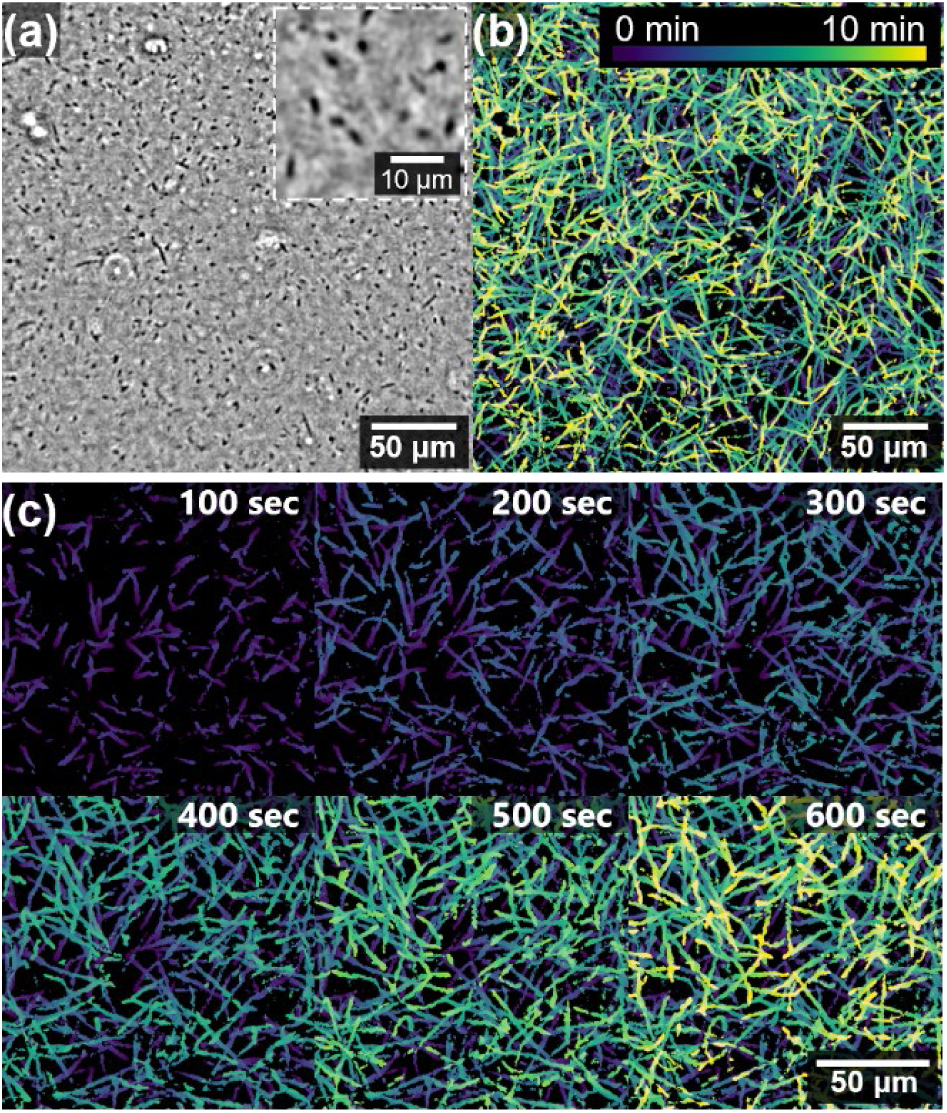
(a) Phase-contrast microscopic image of bacteria during BC hydrogel formation, (b) their trajectories superimposed through time-projection, color-coded to indicate temporal progression, and (c) composite time-lapse images illustrating the process of BC ribbon network formation. The color bar represents the elapsed time.

### Trajectory analysis of bacteria

To elucidate the mechanism of BC structure formation, a quantitative analysis of bacterial trajectories was conducted. Trajectories and velocities were determined by tracking the positions of individual bacteria across frames in time-lapse images (**Figure 4a**). The analysis focused on two commonly studied strains ATCC23769 and ATCC53582. The velocity of bacteria showed a Gaussian distribution with mean ± SD values of 3.2 ± 1.0 µm/min for ATCC23769 and 3.9 ± 1.3 µm/min for ATCC53582 (**Figure 4b**). These values are consistent with previous studies^2,30,31,33–36^, which report that the *Gluconacetobacter* velocities range from 2 to 5 μm/min, suggesting that the motility varies depending on strain and culture conditions such as temperature, medium composition, and substrate.

**Figure 4.**
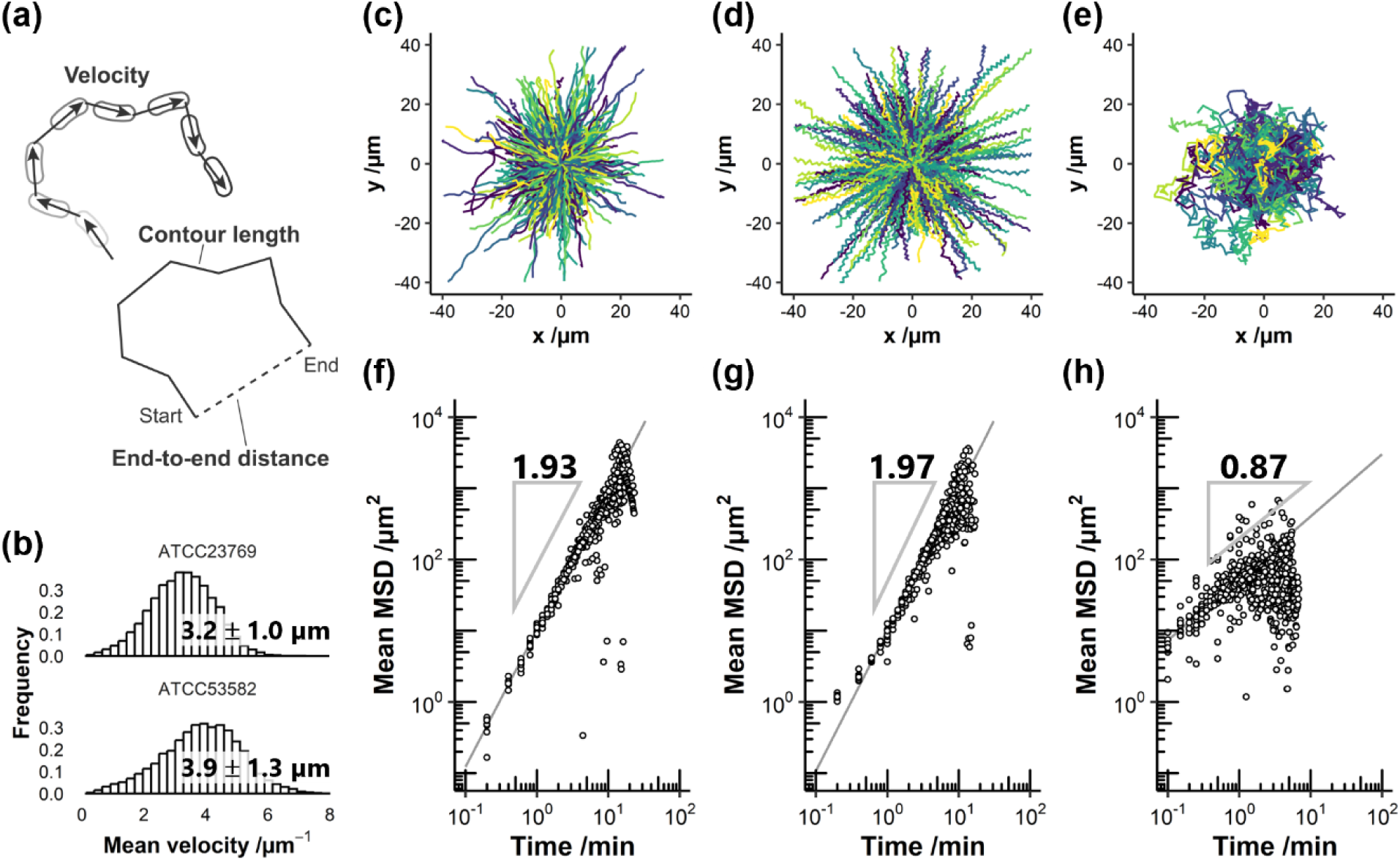
(a) Schematic illustration of bacterial trajectory analysis. (b) Histogram of average bacterial velocities. (c–e) Trajectories of ATCC53582 (c), ATCC23769 (d) and Brownian motion of inactivated bacteria (e). (f–h) Corresponding MSD vs time plots, with fitted slopes indicated.

To clarify intrinsic motility characteristics, the starting points of the trajectories were shifted to the origin and plotted (**Figure 4c–e**). Both strains, ATCC53582 and ATCC23769, exhibited predominantly linear trajectories (**Figure 4c, d**), markedly different from the Brownian motion exhibited by inactivated bacteria (**Figure 4e**).

To further characterize their motility behavior, mean squared displacement (MSD) analysis was performed (**Figure 4f–h**). Bacteria secreting cellulose fibers displayed MSD slopes close to 2 in log–log plots (**Figure 4f, g**), indicative of super-diffusive movement, whereas Brownian-moving bacteria showed a slope near 1 (**Figure 4h**). These results suggest that fiber-secreting bacteria move in a highly linear manner, likely due to the rigidity imparted by the cellulose fiber they produce. Repula *et al*.^31^ also performed an MSD analysis of *Gluconacetobacter* migration, interpreting slopes less than 2 on sub-minute time scales as a cage effect. Consistently, in our ATCC23769 data, a decrease in the slope at short timescales (∼1 min) was observed. We attribute this to the fluctuation in the bacterial center of mass caused by rotational motion as bacteria spiral around their cellulose secretion axis.

### Mechanism of fiber network formation

The formation of BC hydrogels is characterized by the bacterial secretion of cellulose fibers and the subsequent development of a fiber network. While the biosynthesis mechanism of individual BC fibers has been extensively studied, the mechanisms governing the fiber network assembly remain less clear. Because the physical properties of BC hydrogels largely depend on this fiber network architecture^3,40^, elucidating its formation mechanism is essential for advancing material design and performance.

Atomic force microscopy (AFM) revealed that BC ribbons exhibit bundled structures with a right-handed twist (**Figure 5a**), as previously reported^42–46^. Notably, branching and merging points are observed, suggesting that these structures form as fibers bundle and unbundle over time. Time-lapse analysis of bacterial movement identified five distinct behaviors associated with higher-order structure formation (**Figure 5b–f**):

(i) **Tracing**: Bacteria frequently trace pre-existing fibers, moving along the twisted, right-handed cellulose ribbons (the zigzag trajectory prominently shown in **Figure 5c, d**). Multiple bacteria can follow the same trajectory sequentially, indicating that repeated tracing promotes fiber bundling (**Figure 5b**).
(ii) **U-turn**: Collisions with other bacteria or fibers can cause the bacteria to reverse their direction (**Figure 5c**), suggesting bidirectional movement along cellulose nanofibers. This indicates that fibers are oriented with some degree of randomness within the network.
(iii) **Branching by cell division**: Fiber branching occurs as bacteria dividing during secretion separate and continue to produce fibers independently (**Figure 5d**).
(iv) **Branching by physical disturbance**: External physical disruption can cause bacteria to deviate from their original paths, creating branch points (**Figure 5e**). The frequency of such derailments is likely correlated with the bacterial and fiber volume fraction.
(v) **Merging**: When cellulose-secreting bacteria encounter existing fiber tracks, merging points are formed (**Figure 5f**). These events are less frequent compared to branching events.

**Figure 5.**
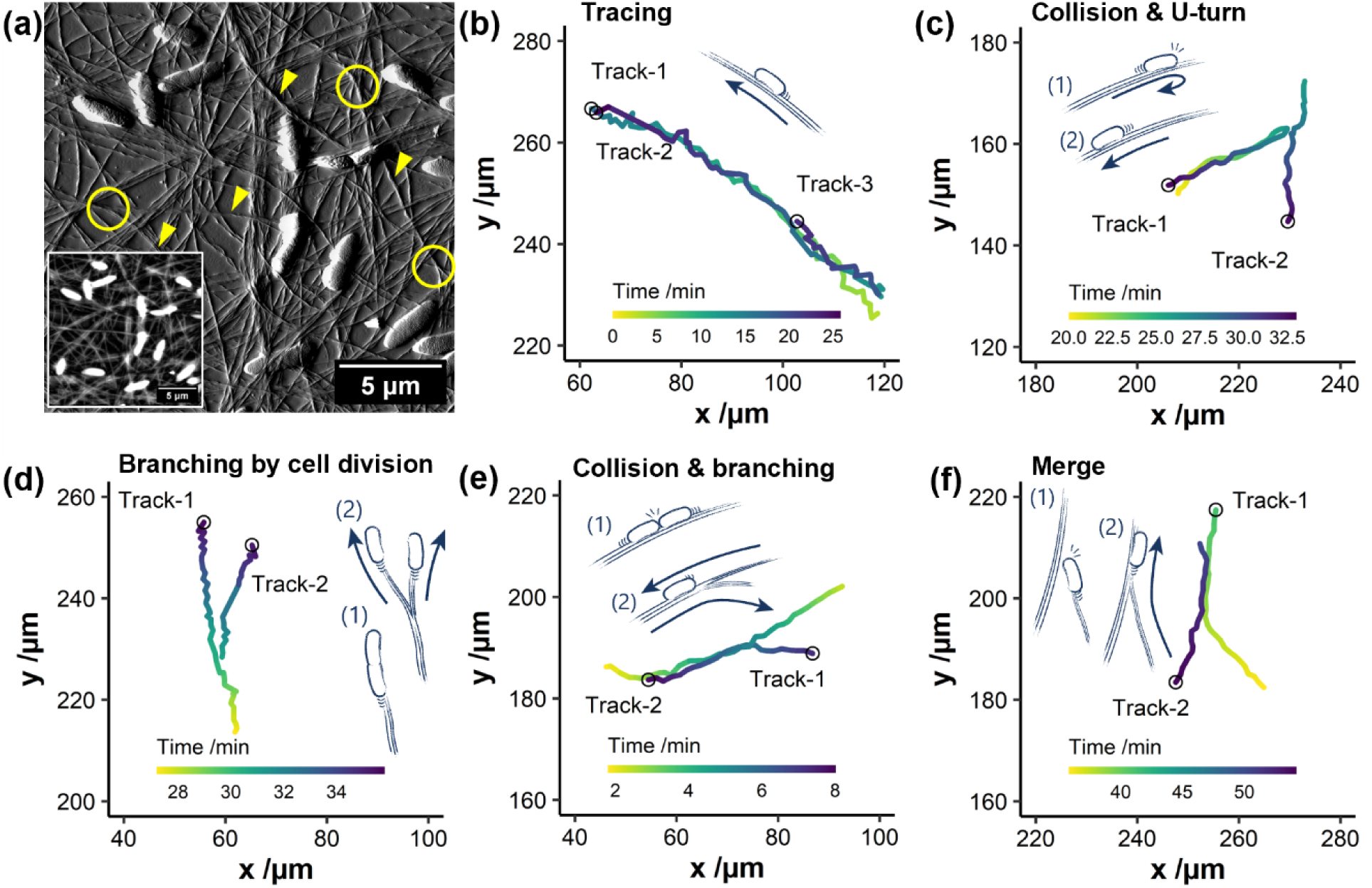
(a) AFM amplitude images of BC ribbons during the initial stage of network formation, with inset height image. Arrowheads and circles denote bundled ribbons and branching points, respectively. (b–f) Representative bacterial trajectories obtained from tracking analysis: (b) tracing, (c) U-turns, (d) branching by cell division, (e) derailment-induced branching, and (f) merging points.

### Formation of nematic domain structures

By culturing *Gluconacetobacter* within a confined space with continuous nutrient and oxygen supply, we successfully induced multi-domain chiral nematic structures (**Figure 6a**). Bacteria within these domains align along a common orientation, as confirmed visually in the inset of **Figure 6a**, while polarized light microscopy confirms the nematic assembly of cellulose fibers (**Figure 6b**). Confocal microscopy further reveals that the structure adopts a chiral nematic (helical) configuration, with the director field rotating continuously with increasing height (**Figure 6c**). The Fourier transform (FFT) image indicates a left-handed helical structure with a half-pitch of ∼18 μm (**Figure 6d**). The emergence of chiral nematic order is attributed to the intrinsic right-handed twisting of individual cellulose fibers.

**Figure 6.**
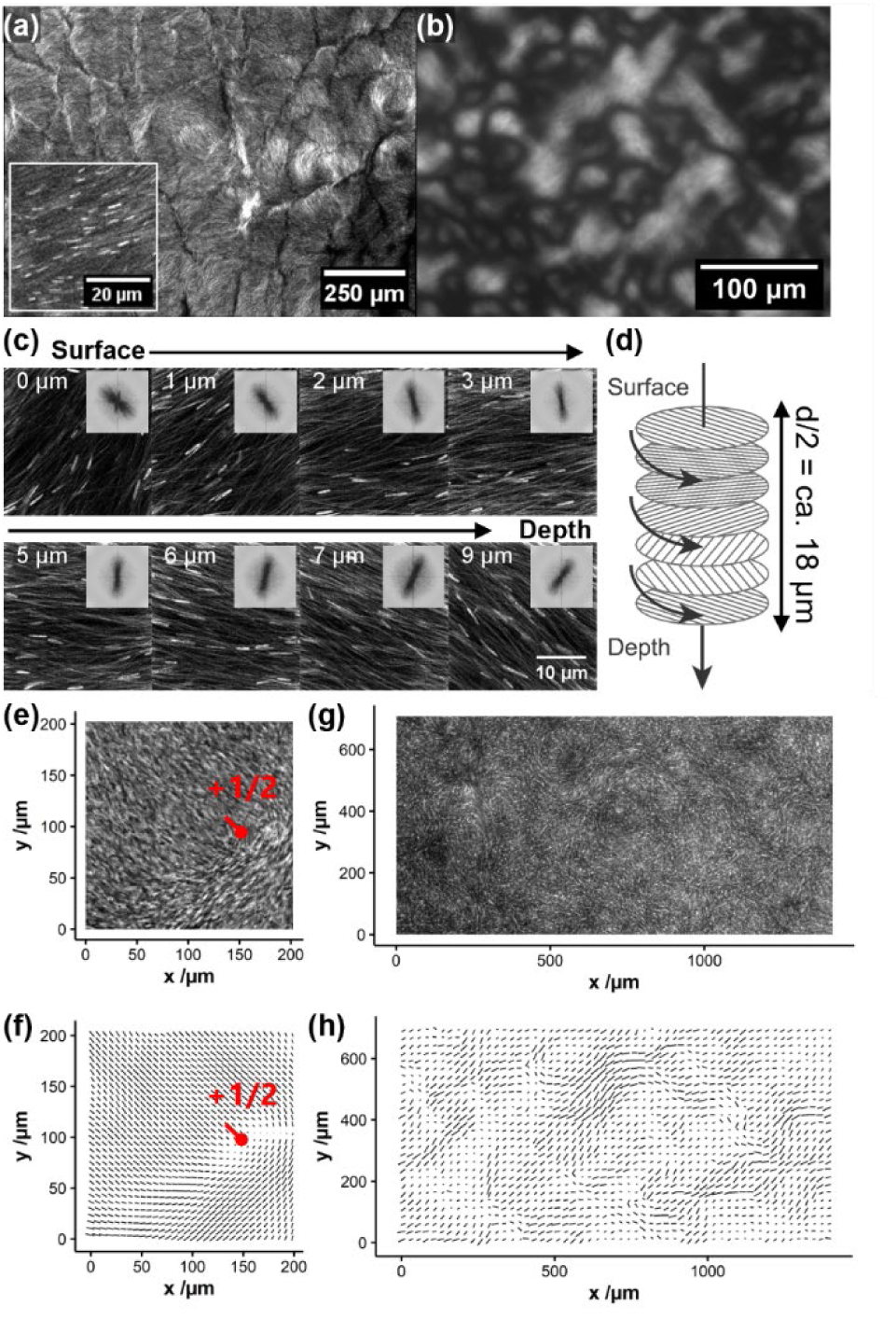
(a) CLSM images showing chiral nematic-like domains in BC hydrogel; the inset depicts bacterial alignment. The orientation of the rod-shaped bacteria is consistent with that of the cellulose fibers. (b) Polarized light microscopy confirms nematic fiber organization. (c) Confocal reconstructions at successive heights with FFT analysis inset, reveals a left-handed helix with a pitch of ∼18 µm (d). (e) Topological defects in phase contrast microscopy; (f) and its corresponding orientation vector map; (g) overview image with width of 1 mm; and (h) vector map.

Orientation analysis of phase contrast microscopic images (**Figure 6e–h**) identified topological defects characteristic of nematic order (**Figure 6e, f**). The domain size is on the order tens of micrometers, indicating that the system exhibits anisotropic properties below this characteristic length scale (**Figure 6g, h**), while appearing macroscopically isotropic due to spatial averaging.

### Time evolution of the nematic phase

Time-lapse observations of the phase transition from isotropic to nematic states indicate that an increase in bacterial and cellulose fiber density triggers spontaneous ordering (**Figure 7**). As bacteria proliferate and continuously secrete cellulose fibers, the effective volume fraction within the chamber increases. During the initial stage (∼0–6 hours), bacterial motion remains disordered; then, around 6.5 hours, loosely ordered structures begin to emerge. These domains continuously grow and merge, and by 10 hours of culture, ordered domains dominate the field of view, indicating a density-dependent phase transition.

**Figure 7.**
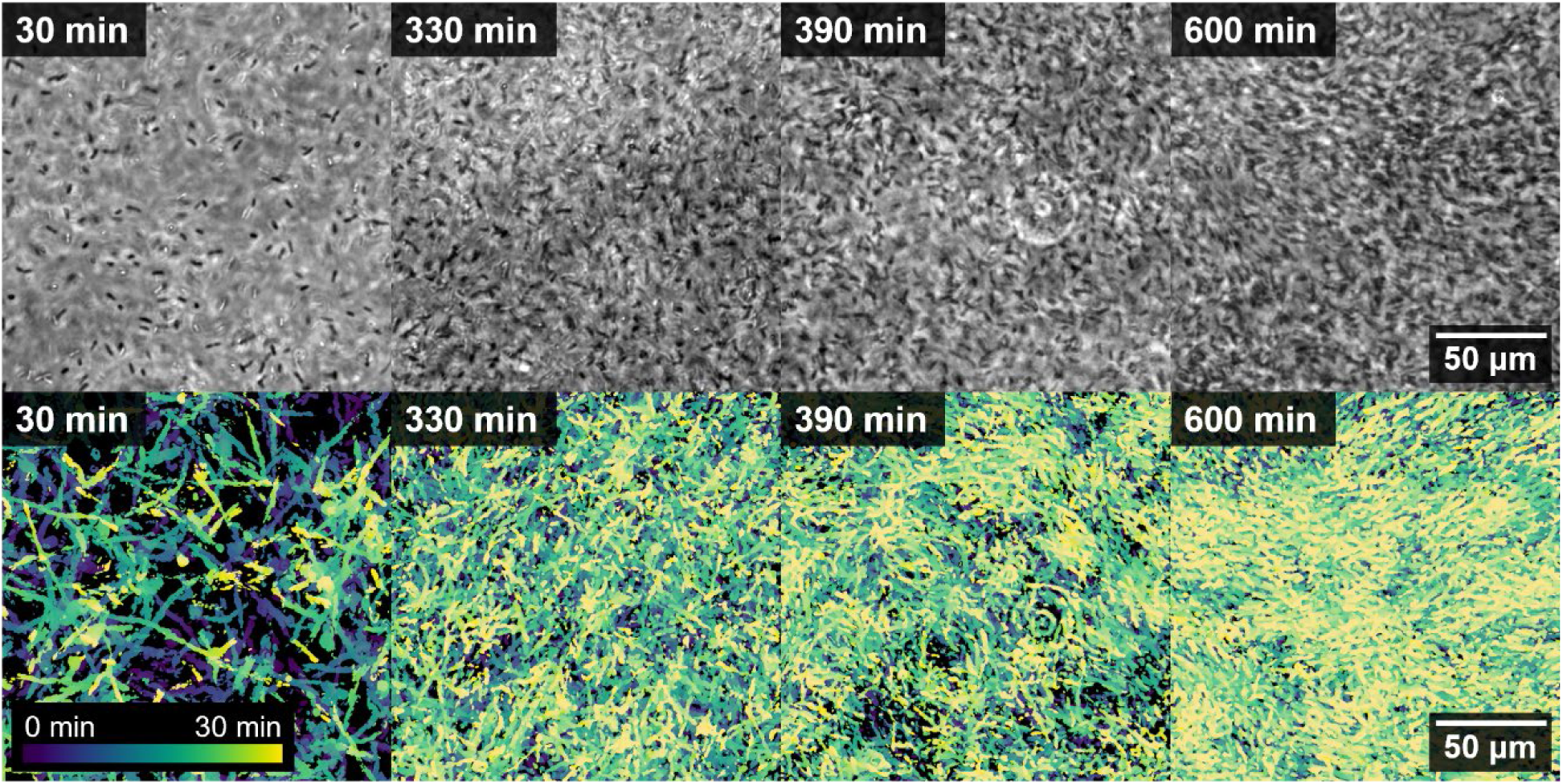
Time-lapse observation of the emergence of the ordered phase during BC hydrogel formation. Bacterial proliferation leads to the gradual development of anisotropic ordered structures from an initially isotropic network. Upper panels present phase-contrast microscopy images illustrating the increase in bacterial density over time. Lower panels display time-projected images (30 minutes duration) of bacterial motility. From approximately 390 minutes onward, loosely oriented domains become apparent, and by 600 minutes, ordered domains dominate the field of view.

### Nematic interactions

In conventional nanocellulose dispersion system, the emergence of nematic or chiral nematic order typically requires relatively high cellulose concentrations^47–50^ (∼10 wt%), consistent with excluded-volume–driven ordering of anisotropic particles. In contrast, the BC hydrogel system exhibits nematic ordering at cellulose concentrations well below 1 wt%, which is unexpectedly low from the perspective of classical theories for rod-like particles, including the Onsager criterion^51^. This discrepancy suggests that nematic ordering in BC hydrogels cannot be explained solely by equilibrium phase behavior of dispersed cellulose fibers. Instead, cellulose fibers secreted by bacteria form an interconnected network whose effective volume fraction increases as bacterial proliferation fills the available space. Moreover, the fibers extend over length scales far exceeding the bacterial size, enabling anisotropic interactions over long correlation lengths, consistent with previously reported liquid-crystalline systems^31,52–55^. These factors, together with the continuous, active deposition of cellulose fibers, likely promote nematic ordering even at low overall cellulose concentrations. A quantitative description of this non-equilibrium ordering mechanism will require further systematic investigation.

### Formation of vortex structures

Occasionally, vortex-like loop structures are observed within BC hydrogels (**Figure 2c**). Confocal microscopy revealed that these loops have diameters of 19 ± 4 µm, several times larger than individual bacteria (**Figure 8a, b**). Once formed, bacteria tend to trace these circular paths repeatedly (**Figure 8c**), reminiscent of swirling behaviors reported previously^30,34,53^. Notably, longer bacteria (length: 12.6 ± 3.7 μm, mean ± SD) more frequently participate in loop formation compared to the general population 3.2 ± 1.5 μm (mean ± SD) (**Figure 8d**). Trajectory analysis indicates that these elongated bacteria often follow wavy, spiral-like paths, likely reflecting helical motion. Such coil-like movement may facilitate the spontaneous closure of paths, leading to vortex loop formation. When multiple vortex loops coexist and interact, the system may develop vortex lattice patterns (**Figure 8e, f**). In contrast to vortex arrays induced by external templates or imposed fields in previous studies^36,56^, these structures arise spontaneously, driven by the intrinsic motility of bacteria coupled with continuous cellulose fiber secretion.

**Figure 8.**
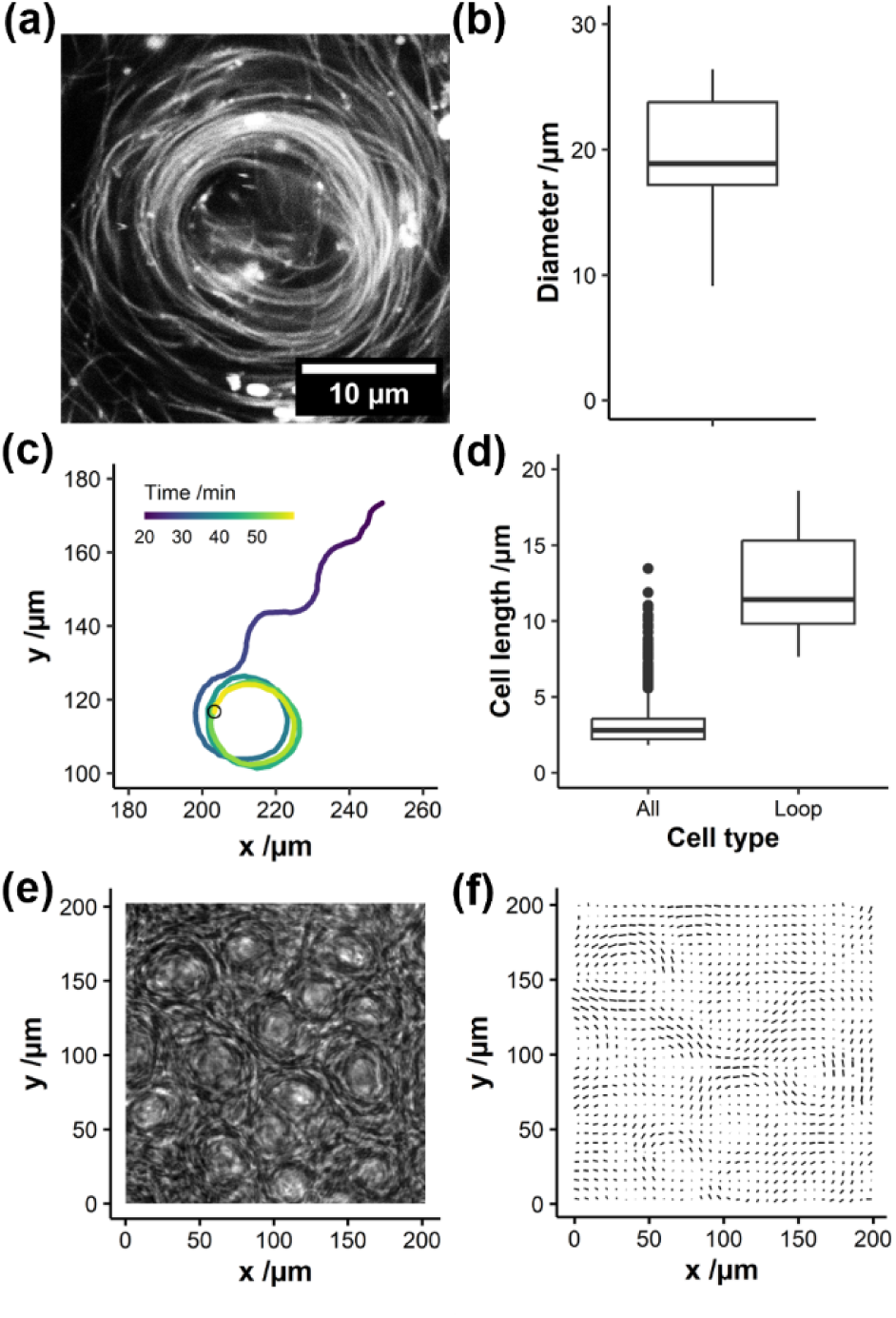
(a) CLSM image of vortex structures in BC hydrogel; (b) diameter distribution of vortex loops. (c) Representative bacterial trajectories forming vortex; (d) boxplot comparing cell lengths of vortex-forming bacteria and the overall population showing preferential involvement of elongated bacteria. (e) Phase contrast image of vortex lattice-like structure and (f) its orientation vector map.

### Model of BC network formation

Based on our experimental observations, we propose a conceptual model for BC fiber network formation (**Figure 9a**). In this model:

i) Bacteria synthesize cellulose fibers and migrate, either creating new paths or tracing pre-existing ones, thereby forming fibrous scaffolds.
ii) Branching points arise when bacteria undergo cell division while actively secreting cellulose fibers.
iii) U-turn behavior occurs upon collision with obstacles, leading to a reversal of bacterial migration direction.
iv) Junction points are generated when bacteria encounter and merge with pre-existing fibers, forming crosslink sites.
v) The interplay of these processes—tracing, U-turns, branching, and merging—gives rise to a crosslinked, mesh-like BC fiber network (**Figure 9a**).

**Figure 9.**
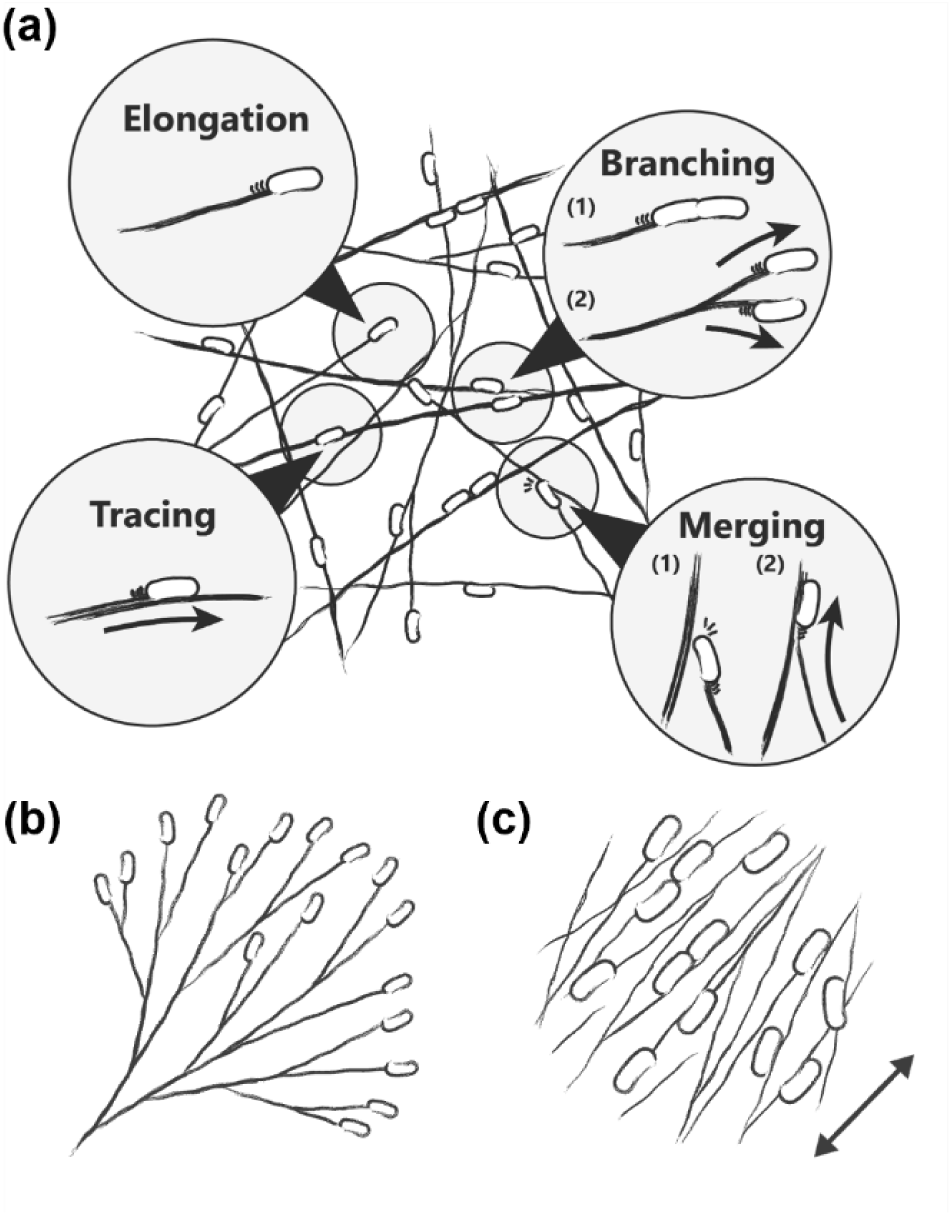
(a) Conceptual model of BC network formation incorporating multiple bacterial behaviors. (b) Tree-like network structure resulting from branching alone; (c) anisotropic, nematic-like network emerging at higher volume fractions.

A simplified model that considers only cell division-driven branching produces tree-like network structures^54,57^ (**Figure 9b**), which are inconsistent with interconnected networks. In contrast, incorporating tracing, U-turns, and merging behaviors results in more realistic, highly interconnected mesh structures.

Furthermore, the transition from isotropic to anisotropic (nematic) BC networks appears to be governed by an increase in fiber and bacterial volume fractions. At higher densities, enhanced bacterial collisions and fiber–fiber interaction, promote nematic ordering and increased crosslink formation (**Figure 9c**).

### Materials design strategy for BC based on spontaneous structure formation

The collective behaviors of cellulose-producing bacteria and the resulting nanofiber architectures provide versatile strategy for tailoring the properties of BC. By controlling key structural features—such as branching frequency, fiber alignment, and vortex formation—it becomes possible to tune the mechanical, optical, and electrical properties of BC materials (**Figure 10**). For example, adjusting the crosslink density directly affects elasticity and toughness, while alignment of nematic domains can induce anisotropic mechanical, electrical, and optical properties. Furthermore, incorporating vortex-like patterns could yield materials with combined extensibility and strength. Realizing such bio-assembled material designs requires a detailed understanding of how active bacterial motion gives rise to emergent BC structures.

**Figure 10.**
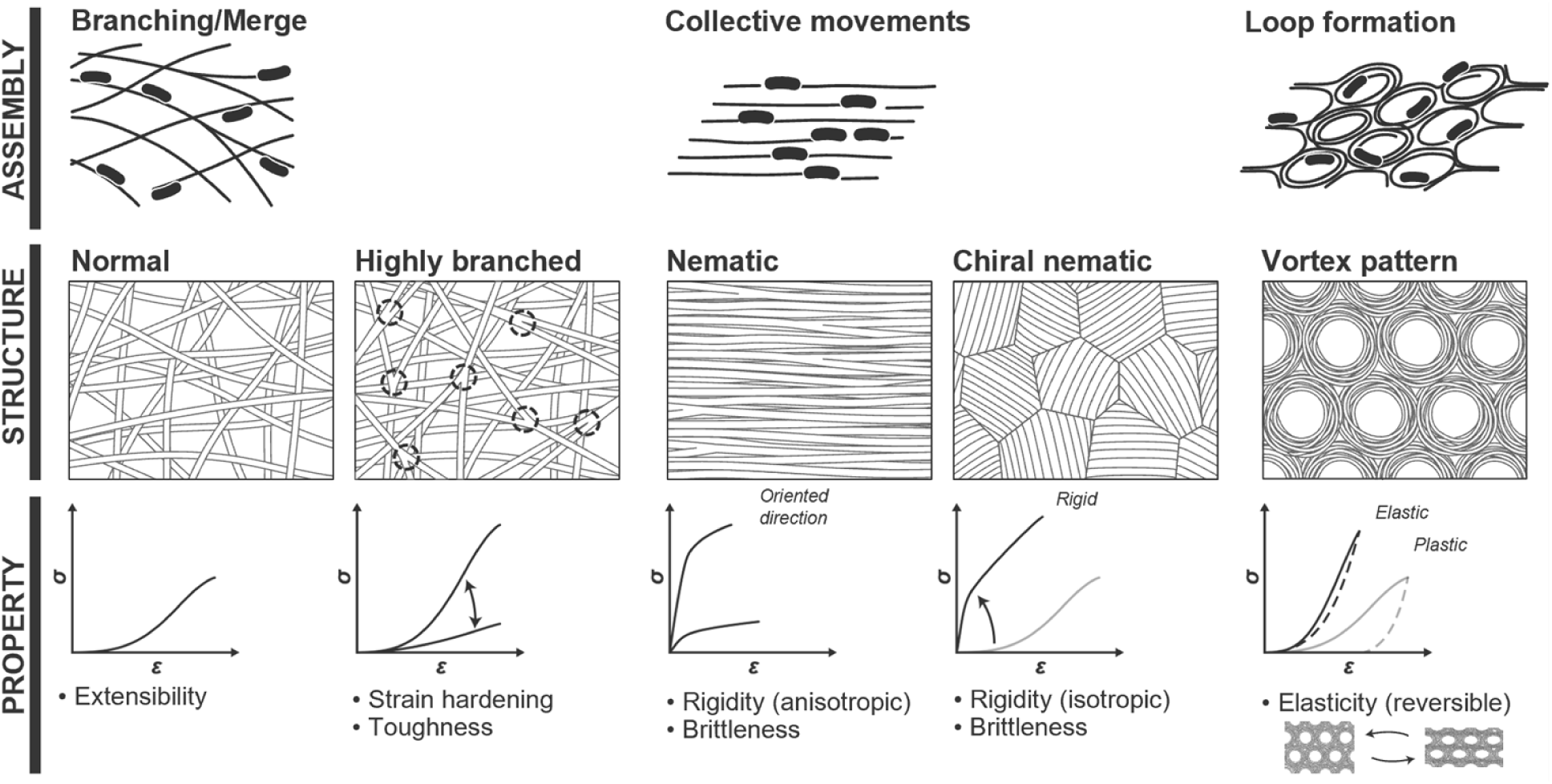
Schematic of bio-assembly-based BC material design: (upper) formation process driven by cellulose-producing bacteria; (middle) resulting structural patterns; (lower) projected mechanical and functional properties.

### Gluconacetobacter as active matter

*Gluconacetobacter*, which propels itself by secreting cellulose fibers, can be regarded as a distinctive type of active matter system. In well-studied swimming bacteria, such as *E. coli* and B. subtilis, hydrodynamic interactions play a crucial role in mediating polar alignment and collective motion^58,59^. By contrast, for *Gluconacetobacter* that migrate by secreting cellulose fibers, hydrodynamic interactions are weak, and nematic interactions become dominant. Moreover, the formation of a crosslinked cellulose fiber network and the use of pre-existing fibers as scaffolds introduce a strong history dependence, whereby past structures constrain and guide subsequent fiber assembly, rendering this system fundamentally different from conventional active matter models. Understanding the collective dynamics of *Gluconacetobacter* from an active matter physics perspective is therefore crucial for elucidating BC network formation and for guiding the rational design of BC-based materials.

## CONCLUSIONS

This study demonstrates that bacterial movement plays a crucial role in governing the formation of BC network architectures, including branching, merging, looping, and orientational ordering. These findings highlight how collective bacterial behaviors directly shape the mesoscale organization of cellulose fibers and, consequently, the physical properties of BC hydrogels. A mechanistic understanding of these bio-assembly processes provides a foundation for tailoring mechanical and functional properties through controlled modulation of bacterial activity and collective dynamics. Ultimately, this work opens new avenues for the rational, bottom-up fabrication of custom BC-based materials by harnessing bacterial motility and cellulose bio-assembly processes.

## Author contributions

G. T. conceived, designed, conducted the experiments, and prepared the draft. T. K. supervised the project, reviewed the draft and edited it into the manuscript. All the authors reviewed the manuscript to be submitted.

## Conflict of Interest

The authors declare no competing financial interest.

## Acknowledgements

CLSM observations were performed using the TCS SP8 at the Center for Advanced Instrumental and Educational Supports, Faculty of Agriculture, Kyushu University, Japan. This work was supported partly by JST SPRING, Grant Number JPMJSP2136 and Grant-in-Aid for JSPS Fellows, Grant Number 25KJ0068.

## Notes

### Competing Interest Statement

The authors have declared no competing interest.

